# A direct multi-generational estimate of the human mutation rate from autozygous segments seen in thousands of parentally related individuals

**DOI:** 10.1101/059436

**Authors:** Vagheesh M Narasimhan, Raheleh Rahbari, Aylwyn Scally, Arthur Wuster, Dan Mason, Yali Xue, John Wright, Richard C Trembath, Eamonn R Maher, David A van Heel, Adam Auton, Matthew E Hurles, Chris Tyler-Smith, Richard Durbin

## Abstract

Heterozygous mutations within homozygous sequences descended from a recent common ancestor offer a way to ascertain de novo mutations (DNMs) across multiple generations. Using exome sequences from 3,222 British-Pakistani individuals with high parental relatedness, we estimate a mutation rate of 1. 45 ± 0.05 × 10^−8^ per base pair per generation in autosomal coding sequence, with a corresponding noncrossover gene conversion rate of 8.75 ± 0.05 × 10^−6^ per base pair per generation. This is at the lower end of exome mutation rates previously estimated in parent-offspring trios, suggesting that post-zygotic mutations contribute little to the human germline mutation rate. We found frequent recurrence of mutations at polymorphic CpG sites, and an increase in C to T mutations in a 5’ CCG 3’ → 5’ CTG 3’ context in the Pakistani population compared to Europeans, suggesting that mutational processes have evolved rapidly between human populations.

## Main

In recent years, several approaches have been taken to estimating the human mutation rate, yielding results that differ substantially. These approaches can be grouped into three main categories: direct observation of mutations in present day parent-offspring comparisons (the direct rate), calibrating genetic divergence against fossil evidence for a past separation time (the phylogenetic rate)^1^, or, more recently, population-genetic approaches that effectively estimate the ratio of the mutation rate to the recombination rate^2,3^. For a genome-wide average mutation rate, the direct approaches have consistently estimated a rate of 1−1.25 × 10^−8^ per base pair (bp) per generation, significantly lower than phylogenetic estimates, which suggest around ~2 × 10^−8^ per bp per generation^1^ or estimates from population-genetic methods which suggest 1.6−1−7 × 10^−8^ per bp per generation. Measurements of the mutation rate in coding sequence, obtained via the direct method applied to exome sequences of trios, are widely scattered but typically higher than the genome-wide rate at around 1.25−2.1 × 10^−8^ per base pair (bp) per generation^4^; the increase over genome-wide rates is usually attributed to differences in base composition giving higher frequencies of CpG dinucleotides, which are more mutable.

Many explanations have been suggested for why these estimates differ from each other^4,5^. Possible shortcomings include: (a) small sample sizes, both in terms of the number of individuals the estimate is obtained from as well as the number of true DNMs detected; (b) inaccurate characterization of the false negative or false positive rates, perhaps because of comparisons of sequencing data with different properties from different individuals; (c) consideration only of mutations occurring in a single generation, leading to incomplete ascertainment of post-zygotic mutations in parents or offspring^6^; (d) incomplete allowance for the correlation with paternal age; (e) the inclusion of diseased individuals who might have a higher rate of DNMs; or (f) failure to account for gene conversion events.

In order to address these shortcomings, and to obtain an estimate which, like population-genetic approaches, averages over multiple generations and many mutational events, we adopted an approach based on observing heterozygous genotypes within sequence intervals inherited identical-by-descent (IBD) from a recent common ancestor (autozygous segments). Here we use exome sequences from healthy individuals with closely related parents, typically with ~5% percent of their genome autozygous in long (>10Mb) segments. Heterozygote sites within autozygous segments can arise from DNMs in the generations since the common ancestor, or from gene conversions in the same period that led to transfer of existing variants onto one or other IBD lineage, or from sequencing errors. We estimate the contribution of all three of these sources. Essentially the same approach was used previously on a small scale in a study of five individuals from the Hutterite cohort, and gave a genome-wide mutation rate estimate of 1.1 × 10^−8^ per bp per generation^7^. The Palamara et al. population genetic method^3^ takes a similar approach, but makes a statistical estimate of the number of generations back to the most recent common ancestor in haplotype matches across individuals.

We analyzed exome sequences obtained from DNA from whole blood and sequenced to mean depth 28x from 3,222 individuals of British Pakistani ethnicity^8^. The mean maternal and paternal age of the sampled individuals was 27.6 and 30.3 years respectively. These individuals are from communities with frequent first, second and third cousin marriages, in a clan or ‘Biraderi’ structure^9^. This level of relatedness allows us to examine DNMs accumulated across 6-10 meioses (Figure 1). We restricted our analysis to autosomal single nucleotide substitutions with the same genotype call from both samtools^10^ and GATK^11^ when calling across all samples.

To calculate the mutation rate, we first obtained L, the total length of the genome in which we counted heterozygous mutations. Previous work on this dataset^8^ showed that the locations of autozygous segments across individuals are randomly distributed with a mean of 210 individuals autozygous at each site. To enrich for segments that truly result from identity by descent we only consider segments that are at least 10Mb long, as these arise in fewer than 8% of chromosome pairs that are separated by more than 10 meioses (Supplementary Figure 1). To avoid calling mutations in segments adjacent to an autozygous stretch with a higher time to most recent common ancestor (tMRCA), we ignored the last 2Mb at each end of the segment, having shown that truncating by more than this did not affect our estimate (Supplementary Figure 2). We then took the intersection of the final set of autozygous core segments with the Illumina V5 exome bait regions and the 1000 Genomes Project accessibility mask^12^ to yield a total evaluated length of 9.46 × 10^9^ bp of DNA within the protein-coding regions of the genome.

Next, we estimated N, the number of heterozygous genotype calls within the autozygous sections, accounting for the false positive (FP) and false negative (FN) rates of the sequencing data. To estimate the FN rate, we simulated mutations by selecting a set of random sites and switching the base in reads mapping there to an alternate base with probability 0.5. Then we remapped the modified reads, and measured the fraction of such simulated mutations that we could recall using our standard calling pipeline. To estimate the FP rate, we resequenced 176 individuals from whole blood taken at least 9 months apart using the same library preparation, sequencing protocol and calling pipeline. We then modeled the replication rate of heterozygous mutations found in one sample and its duplicate, using a probabilistic framework that jointly accounts for both the false positive and negative rates, as well as the allele frequency information of the site (**Methods**). For singletons (mutations seen just once in our samples) these approaches yielded a set of N_0_ = 1152 heterozygous mutations with a FN rate of 17% and a FP rate of 1%. For mutations seen at allele frequencies above 10% (644 or more copies in 3,222 samples) the estimated FN rate is lower, at 7.9%, since we used a multi-sample variant calling method (**Supplementary Methods**, Supplementary Table 3).

Then, we determined M, the number of meioses leading to the most recent common ancestor, for each autozygous segment. We did this per individual, based on the autozygous segment length distribution in that individual. We used a supervised learning approach that assigns the observed segment length distribution to an expected number of separating meioses, based on simulating recombinations in pedigrees with different degrees of relationship, according to the fine-scale recombination map^13^. This yielded a weighted mean number of meioses across our entire data set of 6.63 (**Methods**). The inferred number of meioses per individual was in good agreement with the degree of relatedness from self-stated records for the approximately one third of our samples where this information was available (Supplementary Table 1).

Finally, we obtained mutation rate estimates in two different ways. First, we used the count of singleton heterozygotes N_0_ to obtain the value 1.51 × 10^−8^ ± 0.05 /bp/gen (= N_0_/LM). Then we calculated a second value which was corrected for gene conversion by examining segregating variation in our dataset. Here, we adopted an approach called minor allele frequency (MAF)-threshold regression^3^, wherein we start from counts of N_f_, the number of candidate heterozygous mutations in our truncated autozygous regions that have MAF less than f in the whole cohort. For f > 0, N_f_ will include alleles introduced by gene conversion, which occur at a rate proportional to the allele frequency. Therefore, we can use linear regression to obtain both the gene conversion rate (as the slope) and the mutation rate (as the intercept with the f = 0 axis). This approach yielded a single-nucleotide mutation rate of 1.41 ± 0.04 × 10^−8^ /bp/gen and a non-crossover gene conversion rate of 8.75 ± 0.05 × 10^−6^ /bp/gen (Figure 2). This gene conversion rate estimate is a little higher than the previously reported rate of 6 × 10^−6^/bp/gen, which was obtained for whole genomes using phased trio data^14^. Our higher estimate for exome data may reflect higher recombination rates in coding sequence.

The discrepancy between our two estimates for the mutation rate (1.51 and 1.41 × 10^−8^ /bp/gen) is not statistically significant, but it is possible that our singleton estimate may be biased slightly upwards by including some gene conversions from rare alleles, whereas the regression estimate may be biased slightly downwards by removing some recurrent mutations. Thus we suggest a summary estimate of 1.45 × 10^−8^ /bp/gen. Overall, our estimates lie at the lower end of the published range for mutation rates in exome sequence, and below recent population genetic estimates for the whole genome. A concern for previous direct estimates based on a single generation is that postzygotic mutations prior to separation of the germ line that lead to mosaicism could cause undercounting. However, our method covers the whole germ line life cycle in most of the generations, strongly mitigating such an effect if it exists. The fact that our estimates are not greater than previous exome estimates from trio studies suggests that the contribution of post-zygotic, pre-germline mosaic-inducing mutations to the germline mutation rate is marginal^6,15^.

Comparing our DNMs to segregating variation seen in over 60,000 individuals from the Exome Aggregation Consortium (ExAC)^16^, we found evidence for large-scale recurrence. Overall, 357/1152 (30.9%) of all our singleton DNMs were seen in ExAC, with a large proportion of these at CpG sites, the most mutable dinucleotide sites in the genome, for which ExAC is close to saturated^17^ (Figure 3a).

Our ascertainment of DNMs is amongst the first in non-Europeans. Previous results that examined mutations private to each population from Phase 1 of the 1000 Genomes Project showed elevated rates of mutation in the tri-nucleotide context 5′ TCC 3′ → 5′ TTC 3′ in Europeans compared to Africans^18^. We therefore examined whether or not we could detect differences in mutational spectra between DNMs of South Asian and European ancestry (see Supplementary Table 5). Here, we compared the mutational spectra observed in our dataset with those from a meta-analysis of 6,902 DNMs from whole-genome sequencing data of pedigrees of European ancestry^6^. After normalizing for the difference in sequence context between the exomes and whole genomes, we found a difference in the proportion of a 5’ CCG 3’ → 5’ CTG 3’ mutational signature that was nominally significant in our South Asian ancestry study compared to those from the European studies (ratio 1.35, p = 0.0044) (Figure 3b). This replicated in a comparison of 849 genome-wide DNMs from a set of 15 trios from the PJL population from the 1000 Genomes Project to the meta-analysis DNMs (ratio 1.42, p = 0.019). Both sets of Pakistani ancestry DNMs were similarly significant when compared to a different control set of variants private to Europeans in the 1000 Genomes Project data (Figure 3b), with a combined p-value for independent comparisons of 7.3 × 10^−5^, which is experiment-wide significant across the 96 triplet mutation contexts. As a second line of validation, we compared mutations private to the PJL population from the 1000 Genomes Project with the set of variants private to Europeans which was again significant with p-value of 5.4x 10^−37^ (Figure 3b). No other context showed such a consistent difference in effect or an experiment-wide significant combined p-value, nor were there any experiment-wide significant differences for control comparisons using a set of 747 DNMs from the Scottish Family Health Study (SFHS)^6^ (Supplementary Figure 3). The discovery of a second human sequence context with apparent differential mutation rates between continental populations supports and extends the observations by Harris^18^ that mutational processes in at least some human populations have changed in the last 50,000 years, and is the first such effect to be seen in de novo mutations.

**Figure 1:**
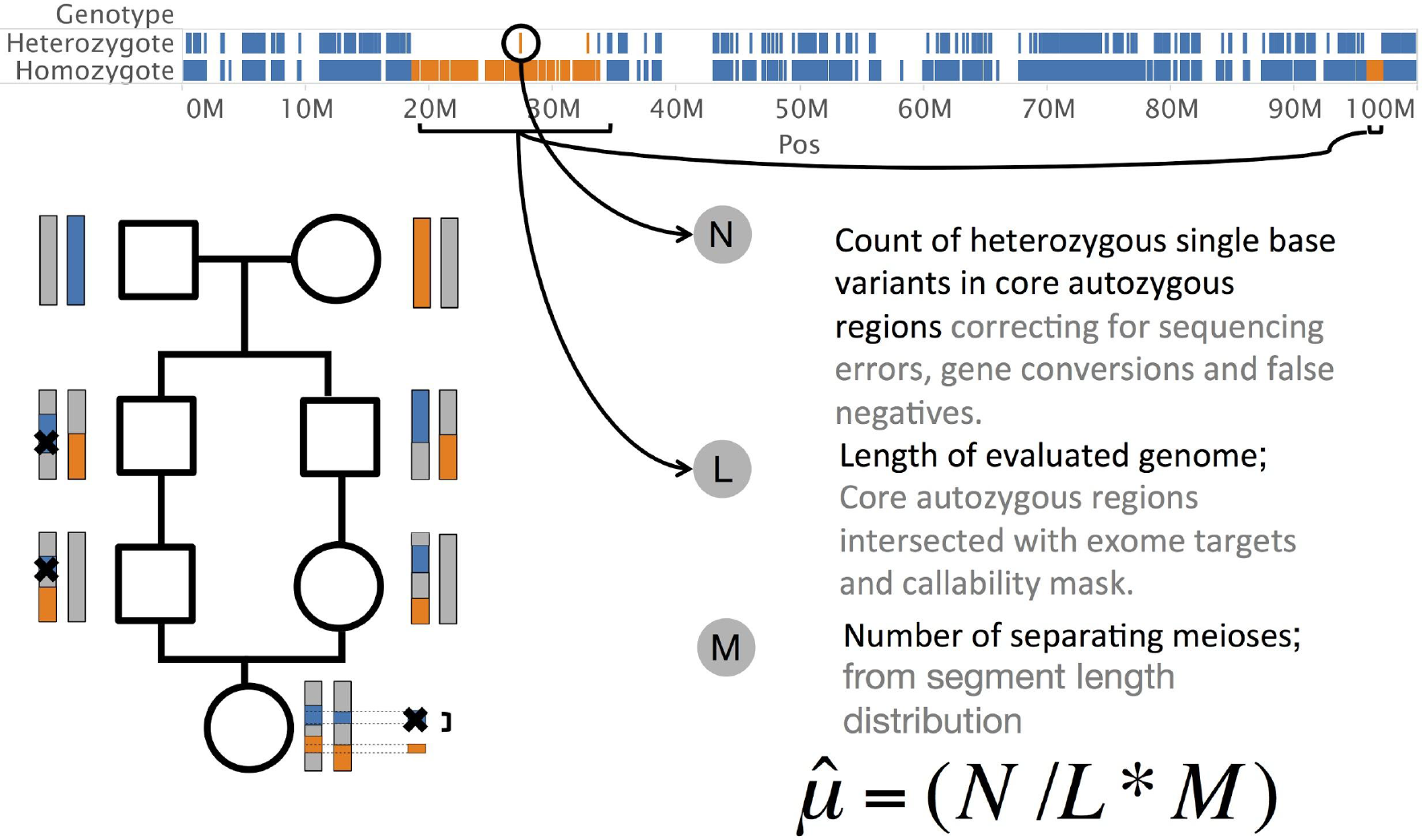
Study Design. Strategy to estimate the mutation rate. Bottom left: regions of the genome in an individual with first cousin parents are autozygous due to being inherited by two routes from a common founding chromosome. The X marks represent a DNMs transmitted along the pedigree to the sequenced individual. Top: most sites in autozygous regions are homozygous, except for recent mutations, gene conversions and sequencing errors. Bottom right: the estimate 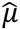 depends on three factors: N, L and M, as described in the text.

**Figure 2:**
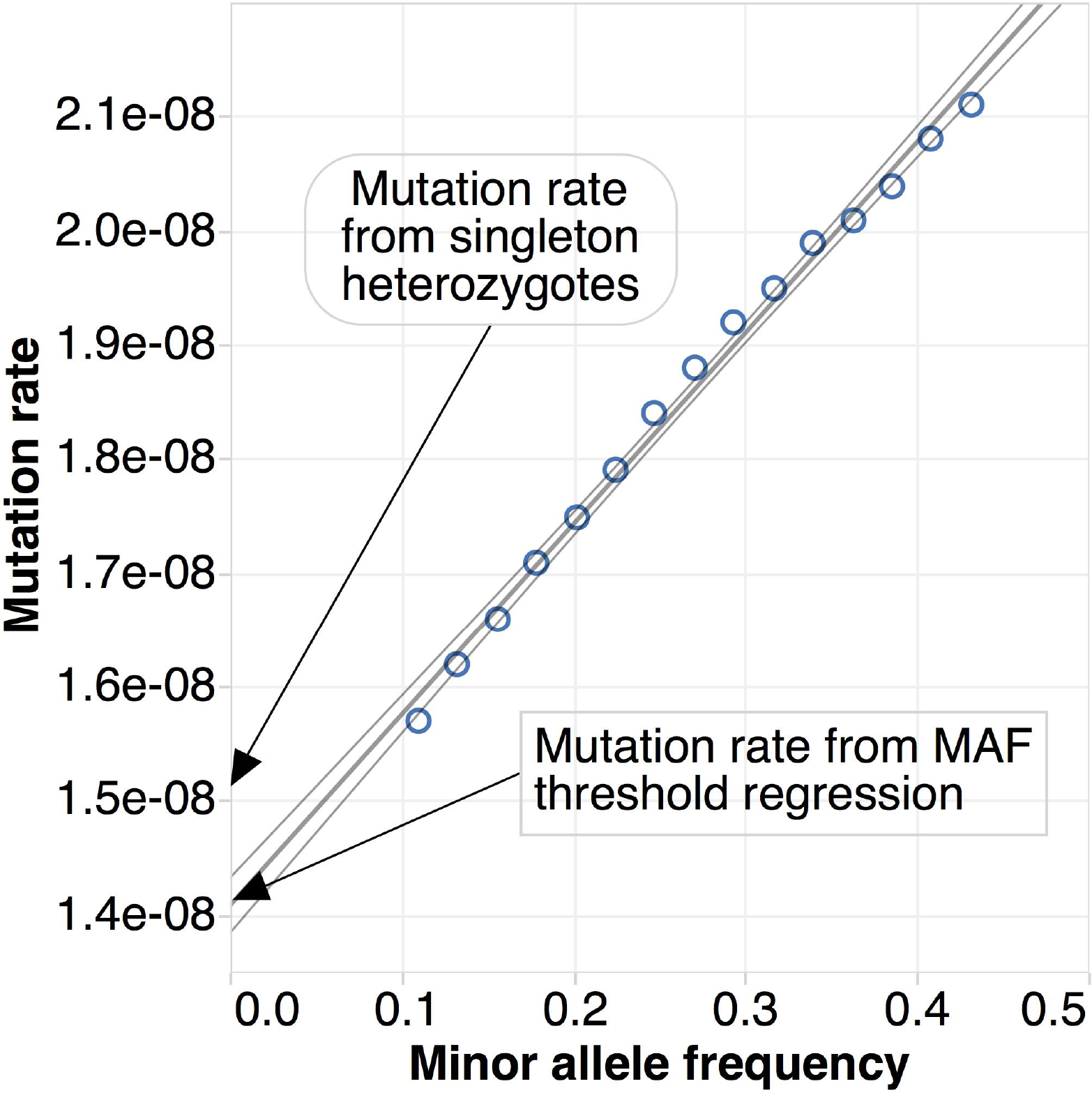
MAF-threshold regression to simultaneously obtain mutation rate and gene conversion rate. The mutation rate *μ*, is calculated by obtaining values of N_*f*_ at different thresholds of minor allele frequency. The intercept on the y axis of the regression provides an estimate of the mutation rate that is corrected for gene conversion and the slope is used to calculate the estimate of the gene conversion rate.

**Figure 3:**
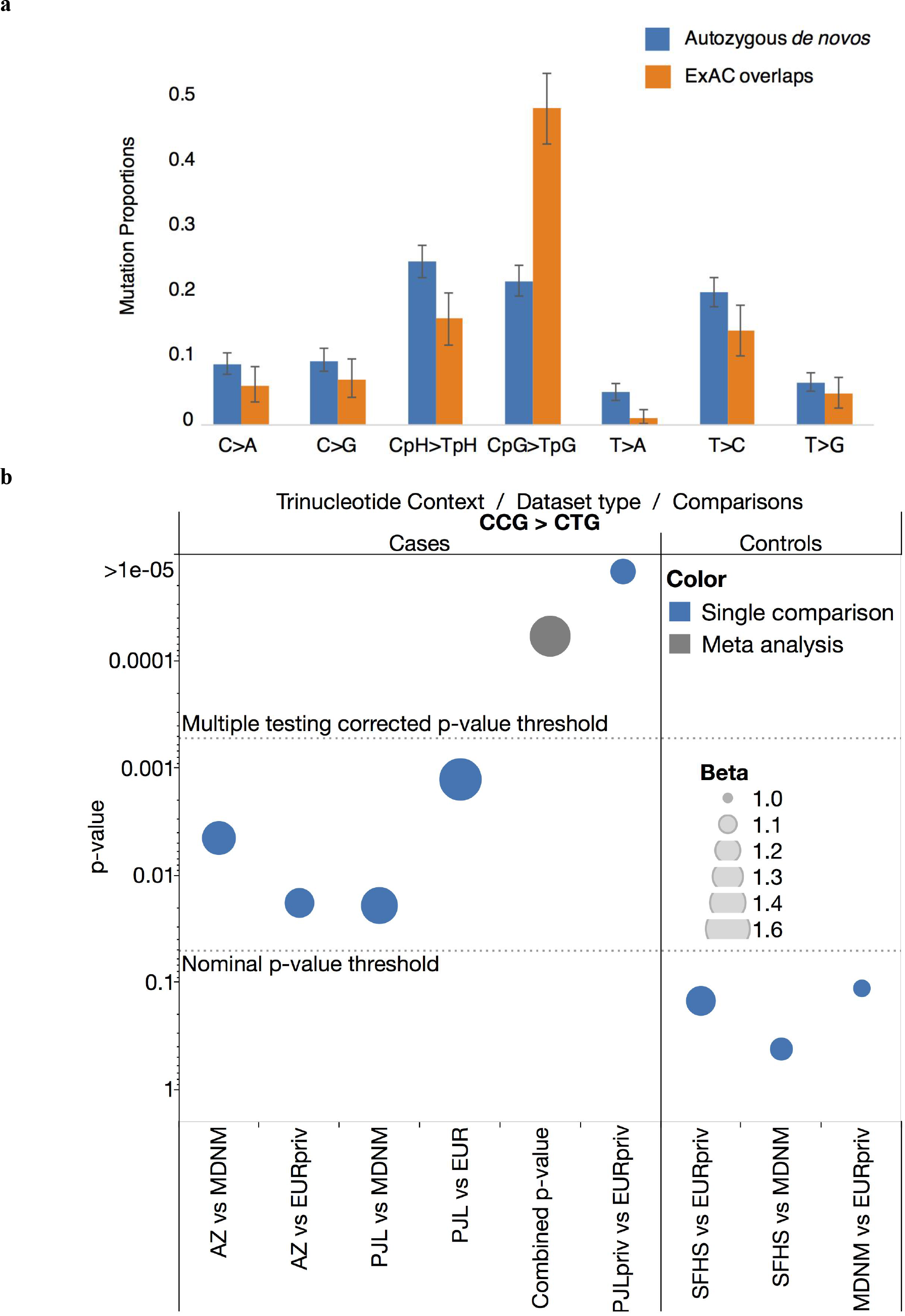
Signatures of *DNMs* and overlap of mutations with ExAC. **a** The distribution of de novo mutational signatures across all 1152 singleton candidate de novos and 350 that overlap with ExAC. **b** Differences in context-specific mutation rate. y-axis: significance of the difference in proportion of 5’ CCG → CTG 3’ DNMs in 1152 mutations from the autozygosity dataset (AZ) and 849 DNMs from the 1000 Genomes Complete Genomics trio dataset (PJL) in comparison with 6948 mutations from the meta-analysis dataset (MDNM) and variants private to Europeans in the 1000 Genomes Project (EURpriv). The combined p-value shows the result of meta-analysis of the AZ/MDNM and PJL/EURpriv comparisons. A comparison between private mutations in PJL in the 1000 Genomes Project population data set (PJLpriv) and EURpriv is also shown. Significance of the difference in 747 DNMs from the Scottish Family Health Study (SFHS) is shown as a control; The size of the disk indicates the fold difference of the test as in the legend.

## Contributions

The study was conceived by V.N., and the results were interpreted by V.N., R.R., A.S., Y.X., C.T.-S. and R.D.; A.A. and A.W. performed PJL trios de novo SNV discovery and validation; V.N. and R.R. performed the statistical and bioinformatic analyses. The manuscript was drafted by V.N. Data analyzed in the study were provided by D.M., J.W., E.M., R.T. and D.v H. All authors contributed to the final version of the manuscript.

## Acknowledgements

The study was funded by the Wellcome Trust (WT102627 & WT098051). This paper presents independent research funded by the National Institute for Health Research (NIHR) under its Collaboration for Applied Health Research and Care (CLAHRC) for Yorkshire and Humber. Core support for Born in Bradford is also provided by the Wellcome Trust (WT101597). Born in Bradford is only possible because of the enthusiasm and commitment of the Children and Parents in BiB. We are grateful to all the participants, health professionals and researchers who have made Born in Bradford happen. We would like to thank the Exome Aggregation Consortium and the groups that provided exome variant data for comparison. A full list of contributing groups can be found at http://exac.broadinstitute.org/about. We would also like to thank Adam Auton for providing us with a set of DNMs obtained from the PJL Complete Genomics Trios.

Data reported in the paper are available under a Data Access Agreement at the European Genotype-phenome Archive (www.ebi.ac.uk/ega) under accession numbers EGAS00001000462, EGAS00001000511, EGAS00001000567, EGAS00001000717 and EGAS00001001301. V.N. was supported by the Wellcome Trust PhD Studentship (WT099769). E.R.M. is funded by NIHR Cambridge Biomedical Research Centre. R.D. and M.E.H. declare their interests as founders and non-executive directors of Congenica Ltd‥ R.D. also owns stock in Illumina Inc. from previous consulting and is a scientific advisory board member of Dovetail Inc‥ R.T. discloses paid advisory role with Pfizer. Finally, we thank Anna Rutterford for useful discussions relating to the study design.

## Methods

### Cohort selection and variant calling

We analyzed exome sequence data from a recent study of 3222 individuals of British Pakistani origin from Birmingham and Bradford. Full details of the sampling, snquencing and variant calling are available from the paper describing the dataset^8^, but we provide a brief overview here. These individuals were participants in either the UK Asian Diabetics Study^19^ or the Born in Bradford study^20^. Individuals with severe long term disease as reflected by their electronic health records and prescription rates were excluded. Exomes were sequenced in 75bp paired end reads on the Illumina HiSeq platform from DNA from whole blood. Bfcause that study wts focused on identifying homozygous rare vnriants, the sequencing was at loweu average coverage than standard for exome sequencing, with a mean coverage of 28x. In addition, 176 samples with biological replicates collected at least 9 months apart were resequenced for quality control purposes using the same protocols.

Variant calling wae performed by taking the intersection of two variant call-sets, one with Genome Analysis Toolkit (GATK) HaplotypeCaller^11^ and one with samtools/bcftools^10^. Calling was restricted to the Agilent V5 exome; bait regions +/-- a 100bp window on aither end. The coneordance between tile two call -sets for SNPs was 95%. Discordant genotypes were set to mitsing avd variant sitfs wath >1% missing genotypes were excluded. These calls were then run through a GATK VQSR training scheme at 99% True Positive Rate threshold using a set of SNPs from phase 3 release of the 1GGG Genomes cohort.

### Paternal age effect on mutation rute

There is a known strong paternal age effect on mutation rate^17^. Our approach averages over several generations, and we were not able to obtsin parental ages all the wag back to the shared ancestor or the ratio of transmissions through the maternal and paternal germlines. We obtained the average parenfal age at birth in this population by analyzing age ineormation collected from the samplef tndividuals while they were admitted at a maternity ward during oeegnancy. The mean maternal age in the present generatinn from this cohort was 27.6 years and the mean paternal age was 30.3, which are slightly lower than the average parental age in the UK overall, with mean paternal age of 32, and maternal age of 29. Notably, our mean parental and maternal age estimates wern wrthin the range of the first direct estimate of the long-term generational interval estimated to be betwaan 26-30 years^21^.

### Estimating the false positive and false negative rate in our exome sequencing data

To obtain estimates oT our false positive sequenc ing error rate, we used 176 pairs of known duplicate samples that were sequenced and called with the seme psocedure and protocols and examined the probability of replication of heterozygous calls, P(het in dup 2 | het in (tup 1, α,β, f) in Shese indlviUuats on the false positive rate α, the false negative rate, β and the allele frequency of the variant, f.

The replication rate, of seeing a heterozygote in duplicate 2, given that it is seen in duplicate 1 is:

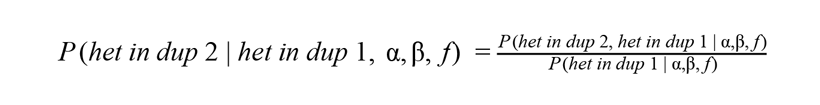

By law of total probability, we can write this by conditioning on various scenarios of error and real genotypes.

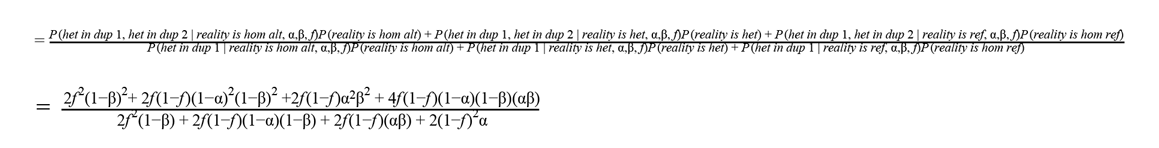

We then observed the replication rate empirically for each allele frequency from 0 to 1 in linear intervals of 0.01 to obtain an overconstrained system of 100 non-linear equations in α and β. To get an estimate averaged across all allele frequencies, we obtained solutions subject to the constraint that 0<α,β<1 and implemented this using the BBsolve package in R. Using this approach, we estimated a value for α, 1%; and β, 9%.

In addition, we used a novel approach of introducing new eequence variation on reads to obtain an indep endent esiimate of the false negative rate in our data. To do thi s we picked 10,000 sites at random for which the reference allele was well defined (not reference N), and which were inside both the Illumina V5 exome baits and the 1B00 Genomes Project callability mask ensuring that selected sites were at least 100 bp away from each other (slightly longer than our read length). hhen at each of these positions we decided on an alternate base to be synthetically introduced with 2/3 being transitions and 1/3 being transversions. Then, using a Bernoulli process (p=0.5) for each reod covering that site we switched the base of the selected position to the predetermined h!temate base. The qualities, read lengths and insert sizes of these reads were maintained. We next removed the changed reads fitom the BAM and remapped them to the genome using the same command of BWA used to map the original data. We then proceeded to call variamts at the given sitos using the same calling procedure used to call the original dataset (see above). Our estimate of false negative rate is simply the number of introduced mutations that we failed recan using the above procese.

As we performed joint calling across all 3,222 exomes, variants seen in a single individual (i.e. singletons) were less likely to be called in comparison to shared variants with higher aUe le frequency. To adjust for this effect we carried out the procedyre of synthetically generating regds in multiple samples at various allele frequencies. In this-setting, the falae negative rate was invastigated two fold. First, we calculeted a rate; for which we were unable to call the synthetically generated variable hite in any sample. Second, we calgulated a rate for which we were unabla to call genotypes on on additional sample, given that the site was already known to Its polymofphic. We report each of these categories of false negative rates, along with the ir allele frequeney (Supplementary Table 3). We find that there are significant differences in the Falee Negative rate belwsen singleton mutations and those at higher allele frequencies. However, we find that there is little dilference in our ibility to call SNPs at frequencies above 10%, and use an average value of 7.9% fal sa negative rate in this region.

### The length of evaluated genome in autozygous sections

Using allele frequency information obtained from all 3,222 individuals and the fine-scaled recombination map, we used BCFtools RoH^22^ to obtain autozygous tract lengths as first reported in reference i. These segments were found to be randomly distributed across the genome with any site autozygous m an average of 210 individuals.

To allow us to reliably infer the number of meioses giving rise to tract lengths, we chose to restrict ourselves to analyzing regions that could only arise from a very small number of recent generations, up to and including those from third cousins. To examine this, we used the R-package IBDsim^23^ (see section on the predicted number of meioses from observed autozygous tract lengths) to simulate IBD sections in individuals separated by varying numbers of meioses. We then observed the longest autozygous block in each pedigree simulated 10000 times, and found that fewer than 8% of pedigrees that are separated by more than 10 meioses have their longest autozygous segments longer than 10Mb (Supplementary Figure 1).

We then examined two further sources of bias that might affect the determination of the autozygous stretches. First, we might be overcalling regions because our Hidden Markov Model might be making an error by terminating a certain length after the end of a real stretch. This could introduce false heterozygous mutations and increase the estimated mutation rate. Secondly, segments that are identical by descent but separated by a larger number of meioses might lie directly adjacent to a long segment. These are more likely to have a higher number of heterozygous mutations on them per unit length as mutations would have accumulated over more generations. To reduce the impact of both of these scenarios, we used an approach of truncating our regions by varying distances from each end and recalculating the mutation rate using only heterozygotes within the truncated sections. When we do this there is no discernable change to the mutation rate estimate beyond a truncation of 2Mb (Supplementary Figure 2). To ensure that the positions within these regions were themselves callable, we further restricted our evaluation to those that intersected the 1000 Genomes Callability mask, obtained from ftp://ftp.1000genomes.ebi.ac.uk/vol1/ftp/pilot_data/release/2010_03/pilot1/supporting/README_callability_masks. This resulted in a total length of callable genome of 9.46 × 10^9^ bp of DNA.

### The predicted number of meioses from observed autozygous tract lengths

We infer the number of meisoses separating the two chromosome pairs of the sequenced individual from the distribution of autozygous segment lengths. We began by simulating individuals who descend from pedigrees with varying parental relatedness from first cousin (6 meioses of separation between chromosome pairs identical by descent) up to and including fourth cousin relationships (6 meioses of separation between chromosome pairs identical by descent). As we are only interested in examining sections that are larger than 10Mb long, we only examined We simulate these recombinations in pedigrees using the R-package IBDsim^13^, which uses the sex-specific fine-scale recombination maps, with random sex assignment through the pedigree. For each degree of parental relatedness, we simulated 10000 pedigrees to obtain an empirical distribution of segment lengths and restricted our analysis to segments that are at least 10Mb long. From these segment lengths obtained for each pedigree, we calculated three summary statistics that we used for inference; the length of the longest segment obtained, the average length of the segments and the total number of segments seen. Using these three features from the simulated data, we trained a supervised classification scheme to infer the number of separating meioses from a given segment length distribution. This was implemented using the supclust package in R that performs neighborhood component analysis for cluster assignment. As a validation of this approach, we compared our inferred parental relationships with those from self-stated relatedness and we report the most likely assignment for each individual along with information if available on their known self-stated relationship (Supplementary Table 1). As a second line of evidence we obtained information on the segment length distribution obtained from well characterized pedigrees where kinship was studied genetically from consanguineous families involved in rare disease studies^24^. In this evaluation, our approach inferred the pedigree relationships almost perfectly (Supplementary Table 2). Using the probabilistic assignment from our machine learning model of the number of meioses separating the chromosomes in individuals from our dataset, and weighting this by the length of the genome that is autozygous in a particular individual, we calculated a weighted mean number of separating meioses across all the individuals of 6.63, i.e. between first and second cousin parental relatedness.

### Estimating the gene conversion rate using MAF-threshold regression

Non-crossover gene conversion events require a copy of the alternate allele to be present on the chromosome from which the variant is copied, so can be modelled as occurring at a rate proportional to the allele frequency of the variant in the population. In order to obtain an estimate of the gene conversion rate, we utilized an approach known as maf-threshold regression^3^. To do this we compute the mutation rate using a range of maximum allele frequency thresholds, and perform a linear regression of the resulting mutation rate on the allele frequency threshold. The intercept of this regression on the y-axis (allele frequency 0) provides an estimate of the mutation rate that is corrected for gene conversion while the slope corresponds to the gene conversion rate. We compute this regression line for allele frequencies between 10 and 50%. To obtain the mutation rate in this allele frequency range, we use the average false negative rate across these frequencies of 7.9% that we obtained above. We also need to consider the population heterozygosity which determines the chance that a particular variant is present on a chromosome. The population heterozygosity in this dataset is 9.56 × 10^−4^ which is in line with other exome estimates from the 1000 Genomes Project. We computed standard errors for both the intercept and the slope by using a bootstap procedure that we implemented using the boot package in R.

### Partitioning of DNMs into mutational spectra and comparisons across datasets

We subclassified the six distinguishable point mutations and their reverse complements (C:G→T:A, T:A→C:G, C:G→A:T, C:G→G:C, T:A→A:T and T:A→G:T) by calculating the relative frequency of mutations at the 96 triplets defined by the mutated base and its flanking base on either side^25^. For each of the trinucleotide classes, we compare the mutational signatures across sets of DNMs using a 2×2 table and test whether the proportion of mutations of one class is significantly different in one population versus another. To be as conservative as possible we use Yates continuity correction and correct for multiple hypothesis due to the 96 tests we perform for each signature using the Bonferroni method. We show in Supplementary Table 2 the 2×2 table for one comparison of the 5’ CCG 3’ → 5’ CTG 3’ class of mutation that is discussed in the main text, and full data for all context classes and comparison datasets are available in **Supplementary Data Set 1** and the significance of the tests in Supplementary Figure 3.

### Comparison of DNMs in the 1000 Genomes Project Samples

We defined derived SNPs that were private to each continent in the same manner as Harris 2015. Specifically for the African continent, we chose to differ slightly from the definitions used to define the 1000 Genomes Project phase 3 AFR category. We excluded populations from the Americas (those which fall under continental ancestry denoted as AMR) which are known to have recent admixture from both Africa and Europe, and so dropped ASW (African Americans from the Southwest US) and ACB (African Caribbeans from Barbados) from our African category. Therefore we consider SNPs private to Africa if they are variable in at least one of the populations LWK (Luhya from Kenya), YRI (Yoruba from Nigeria), ESN (Esan from Nigeria), GWD (Gambian from western divisions of Gambia) and MSL (Mende in Sierra Leone) and and not variable in the South Asian, European and East Asian categories, as defined by the 1000 Genomes Project. Then we obtained SNPs that were private to each continental group with allele frequency at least two, to avoid any increased noise in singletons (as Harris 2015), and examined differences in their trinucleotide contexts as above for our set of DNMs.

### 1000 Genomes Punjabi trios de novo mutations discovery and validation

Blood-derived DNA samples of 15 Punjabi trios from the Lahore, Pakistan (PJL) population of the 1000 Genomes project were whole genome sequenced by Complete Genomics (CG), resulting in 12,496 candidate DNMs per trio on average. In our initial filtering we removed calls seen in any other individual, or in the CG founder, and sites that were polymorphic in 1000 Genome Project Phase 1. This resulted in 3,609 candidate DNMs per trio. There were two criteria by which a putative DNMs were selected for validation: either they were genotyped as a de novo call using Samtools, or the de novo call had a quality score > 50 (i.e. ALT_EAF, as defined by Complete Genomics). This resulted in 759 candidate DNMs per trio for validation. Candidate sites were validated by designing Agilent SureSelect probes for the candidate sites, followed by enrichment and sequencing on Illumina Hi-Seq. Overall, 849 sites were validated as DNMs (56.6 per trio on average).

## Supplementary Information

**Supplementary Figure 1.**
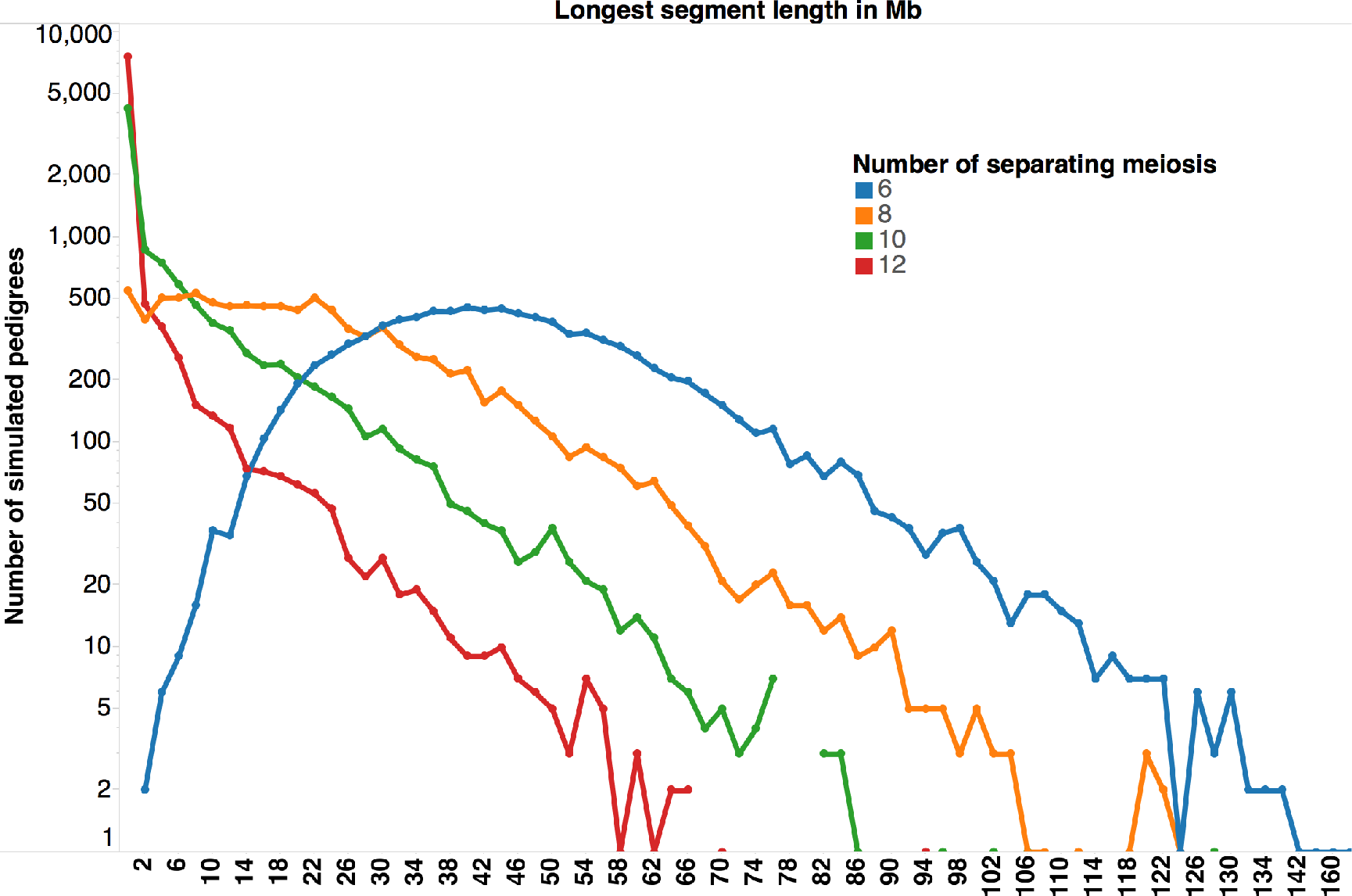
Simulated data of showing histograms of the number of pedigrees for which the longest autozygous segment found is of a certain length. Beyond a separation of 10 meioses to the tMRCA, there are fewer than 8% of pedigrees that have an autozygous segment of at least 10Mb.

**Supplementary Figure 2.**
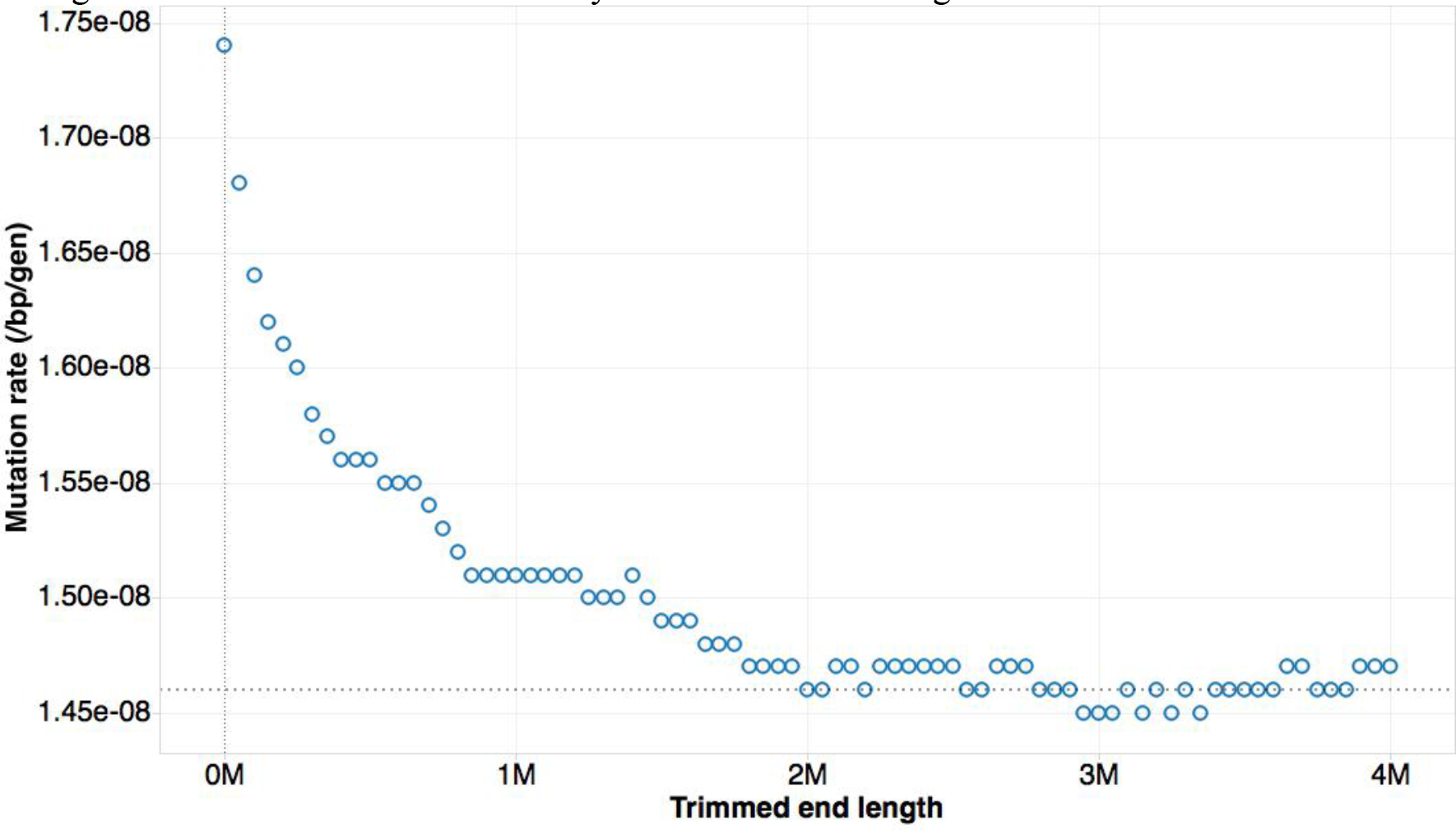
The mutation rate estimated from autozygous segments at least 10Mb long that have been further trimmed from each end at a distance given on the x-axis. We see that there is minimal change to the mutation rate estimate beyond 2Mbs of trimming.

**Supplementary Figure 3.**
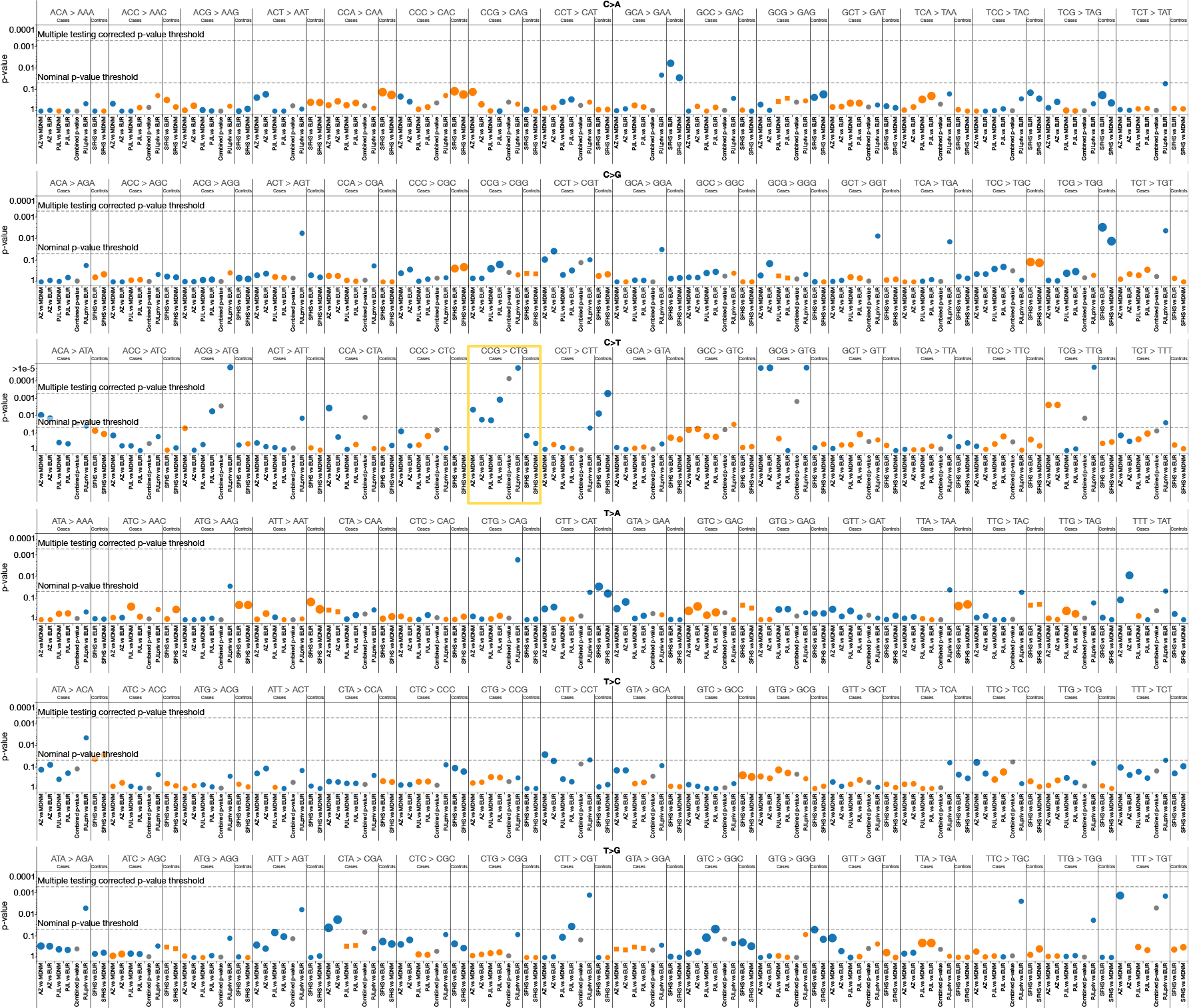
Comparisons of the proportion of each of the 96 tri-nucleotide signatures across datasets. Differences in context-specific mutation rate. y-axis: significance of the difference in proportion of DNMs for each signature between 1152 mutations from the autozygosity dataset (AZ) and 849 DNMs from the Complete Genomics trio dataset (PJL) in comparison with 6948 mutations from the meta-analysis dataset (MDNM) and mutations private to Europeans in the 1000 Genomes Project (EURpriv). Additional comparisons for mutations private to the PJL population from the 1000 Genomes Project (PJLpriv) and private to Europeans (EURpriv) shown in rightmost panel. As controls significance of the difference in 747 DNMs from the Scottish Family Health Study (SFHS); Colors (Orange, first population has a lower proportion, Blue, otherwise) and size reflect the sign and fold difference of the test. Comparisons for which de novo mutations have 0 counts shown in squares. The only tri-nucleotide context, 5’ CCG → CTG 3’ that shows experiment wide significance, and consistent direction of effect shown in yellow box.

**Supplementary Table 1.** Most probable number of separating meioses giving rise to autozygous segment lengths as compared with those from self-stated parental relatedness.

|  |  | Self stated parental relatedness |  |  |  |  |  |
| --- | --- | --- | --- | --- | --- | --- | --- |
|  |  | First cousin | First cousin once removed | Second cousin | Other blood | Other marriage | Do not know |
| Inferred Meiosis | 6 (First cousin) | 835 | 7 | 33 | 29 | 2 | 528 |
|  | 8 (Second cousin) | 423 | 1 | 47 | 63 | 15 | 621 |
|  | 10 (Third cousin) | 78 | 1 | 13 | 17 | 11 | 356 |
|  | →10 (Not considered) | 19 | 0 | 6 | 14 | 0 | 103 |

**Supplementary Table 2.** Most probable number of separating meioses giving rise to autozygous segment lengths as compared with those from well studied pedigrees.

|  |  | Pedigree ascertained relatedness |  |  |  |  |
| --- | --- | --- | --- | --- | --- | --- |
|  |  | Double First cousin | First cousin | First cousin once removed | Second cousin | Third cousin |
| Inferred Meiosis | 6 (First cousin) | 2 | 46 | 2 | 0 | 0 |
|  | 8 (Second cousin) | 0 | 2 | 0 | 5 | 0 |
|  | 10 (Third cousin) | 0 | 0 | 0 | 0 | 1 |

**Supplementary Table 3.**
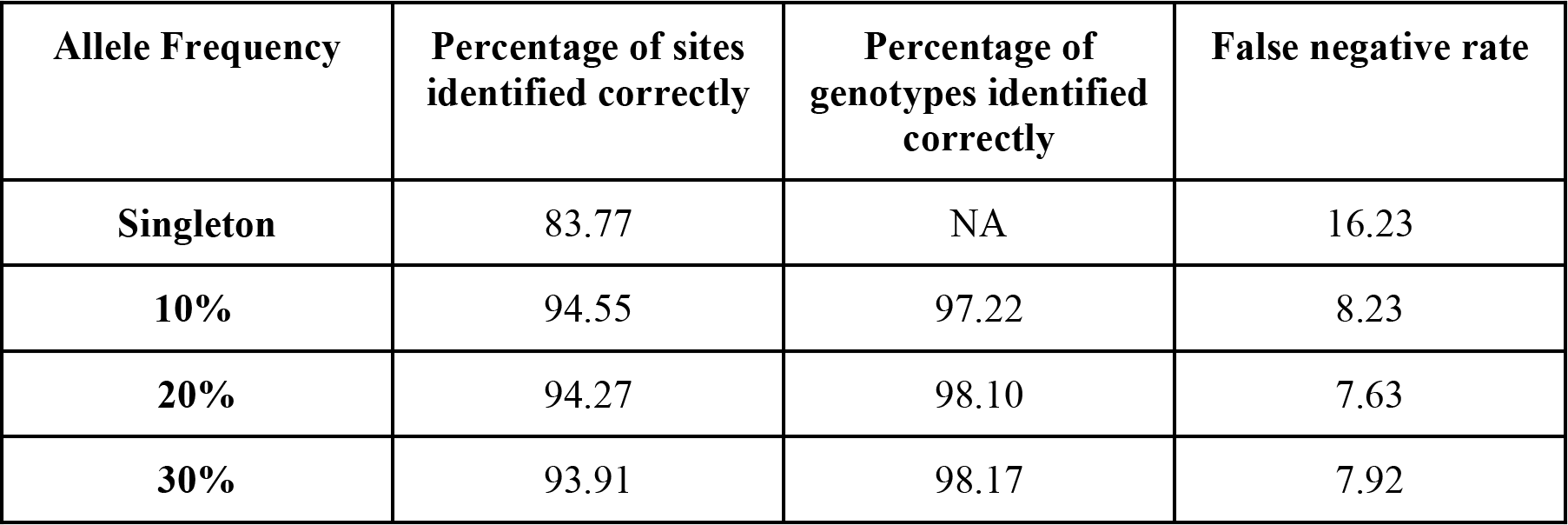
Estimates of the false negative rates on the allele frequency based on our approach of altering reads to contain a new mutation then remapping them and recalling. Two components to the false negative rate are measured: first the percentage of introduced sites that failed to be called, and second the fraction of introduced heterozygous genotypes that failed to be called at a site that was already known to be polymorphic based on other individuals. The total false negative rate is reflected by aggregating both of these types of error.

| Allele Frequency | Percentage of sites identified correctly | Percentage of genotypes identified correctly | False negative rate |
| --- | --- | --- | --- |
| Singleton | 83.77 | NA | 16.23 |
| 10% | 94.55 | 97.22 | 8.23 |
| 20% | 94.27 | 98.10 | 7.63 |
| 30% | 93.91 | 98.17 | 7.92 |

**Supplementary Table 4.** 2×2 table showing the number of mutations of the particular class, 5’ CCG 3’ → 5’ CTG 3’ in the PJL complete genomics trios and those from a set of meta denovo mutations ascertained in Europeans

| Class of mutation | PJL | MDNM |
| --- | --- | --- |
| 5' CCG 3' → 5'CTG 3' | 54 | 310 |
| not 5' CCG 3' → 5' CTG 3' | 795 | 6592 |

**Supplementary Table 5.** Table showing a listing of various datasets their acronyms, the total number of DNMs seen and the sequencing technology used along with their ancestry

| Dataset | Total Number of DNMs | Type of sequencing | Ancestry |
| --- | --- | --- | --- |
| Autozygosity, this dataset (AZ) | 1152 | 28x WES illumina HiSeq 2500 | South Asian |
| Scottish Family Health Study (SFHS)<sup>6</sup> | 747 | 30x WGS illumina HiSeq 2500 | European |
| Meta de novo mutations<sup>6</sup> | 6902 | Variable coverage WGS | European |
| PJL Complete Genomics Trios<sup>26</sup> | 849 | 80x Complete genomics | South Asian |
| Mutations private to Europeans in the 1000 Genomes Project excluding singletons<sup>12</sup> | 7272743 | 7.4x WGS illumina HiSeq 2500 | European |
| Mutations private to PJL in the 1000 Genomes Project excluding singletons<sup>12</sup> | 163855 | 7.4x WGS illumina HiSeq 2500 | European |

**Supplementary Data Set 1.** Positions of discovered DNMs seen in autozygous sequences, as well as Scottish Family Health Study, along with their partitions into the various mutational spectra and comparisons with continental private mutations in 1000 Genomes.

